## Supplementary material for "Perceptual clustering in auditory streaming"

### Supporting information - Graphical model

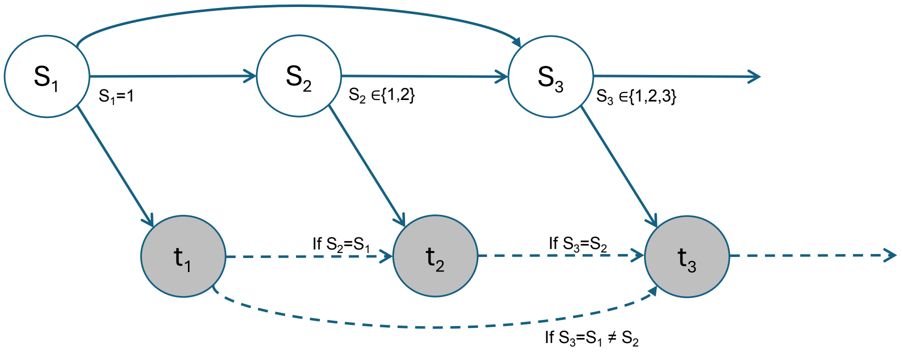


Figure 1: Graphical representation of the main model. Sources (S) generate observed tones (t). First source is always indexed as 1, subsequent sources can either be same or different, with probabilities generated according to the Chinese Restaurant process (see main text Eqs.2-3). The properties of each tone (frequency, on-time, off-time) are conditional upon the previous tone with the same source. For more details see the main text.

### Supporting information - All data and model fits - Exp 1

Supporting figure 2 shows the experimental results from Exp 1 and samples from the model after fitting to the data. Note that a subsection of the data and simulations were presented in the main paper.

Exp 1
Data &
Model

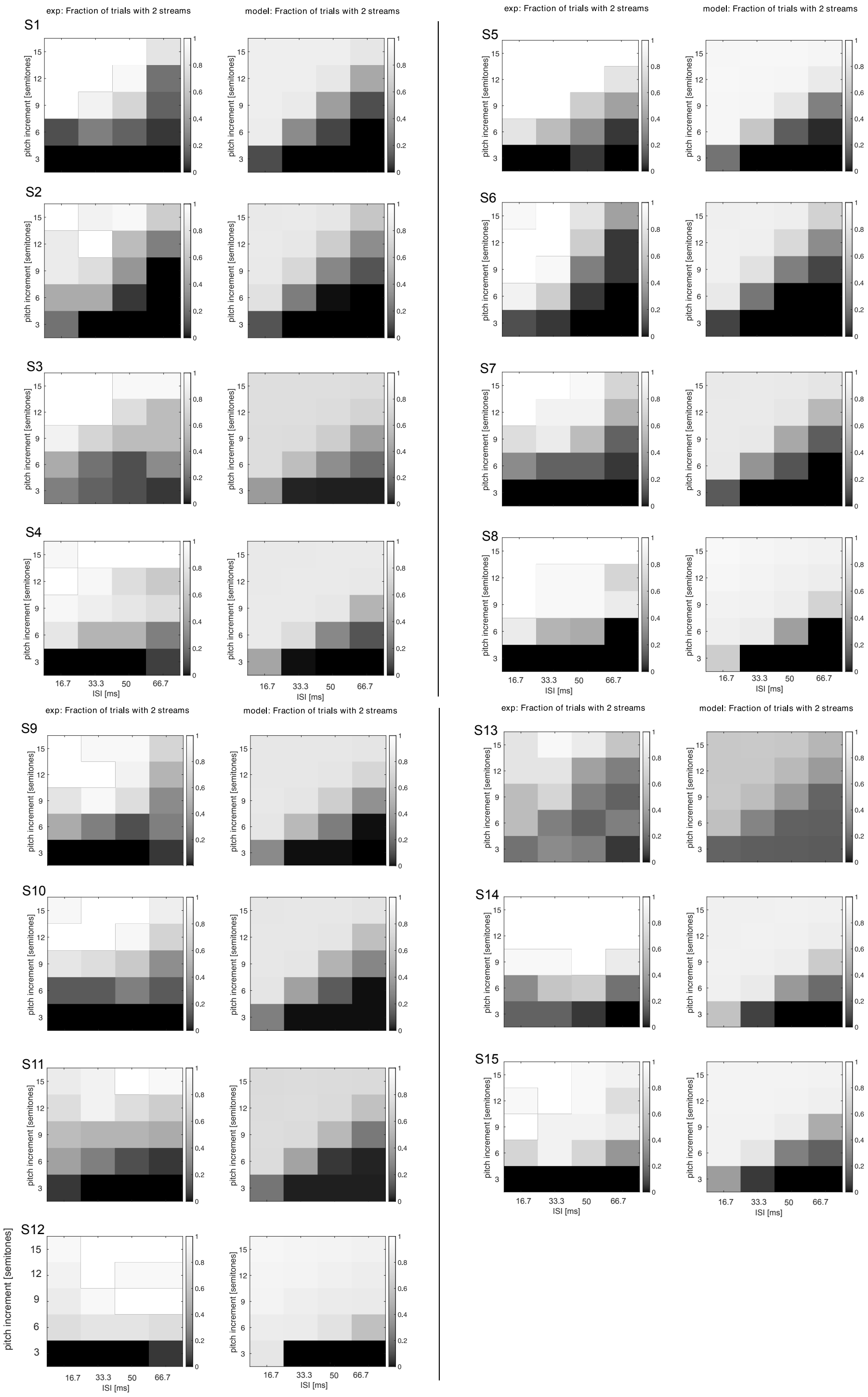


Exp 1
Data &
Model

Figure 2: All data and model fits for Exp 1. Upper left corner shows the subject ID (S1-S15). Experimental data is to the left, reproduction after model fit on the right.

### Supporting information - Parameter recovery

When fitting parameters of a model to a data set it is important to first ensure that the model is specific enough that it is possible to recover its generating parameters when fitting to simulated data sampled from the model.

#### Parameter recovery for Experiment 1 fitting

Parameter values were sampled from prior distributions over the generating parameters $P(onesource)$, $\sigma$ and $\beta$. Based on this parameter set, subject responses were simulated from the model in accordance with how subject were recorded (i.e. same number of trials, same number of conditions). Given this simulated data set (for which we know the true generating parameters) the model was fit unto, using the same method as for fitting to the real subject data. The fitting on simulated data reveals a set of fitting parameters that can then be compared with the generating parameters. Figure 3 shows the recovered parameters for $P(onesource)$, $\sigma$, and $\beta$, plotted against the original generating parameters. The fitted parameters are reasonably close, implying that the model is correctly specified. Obviously, more trials per condition would decrease noise and therefore lead to better fits of the parameters.

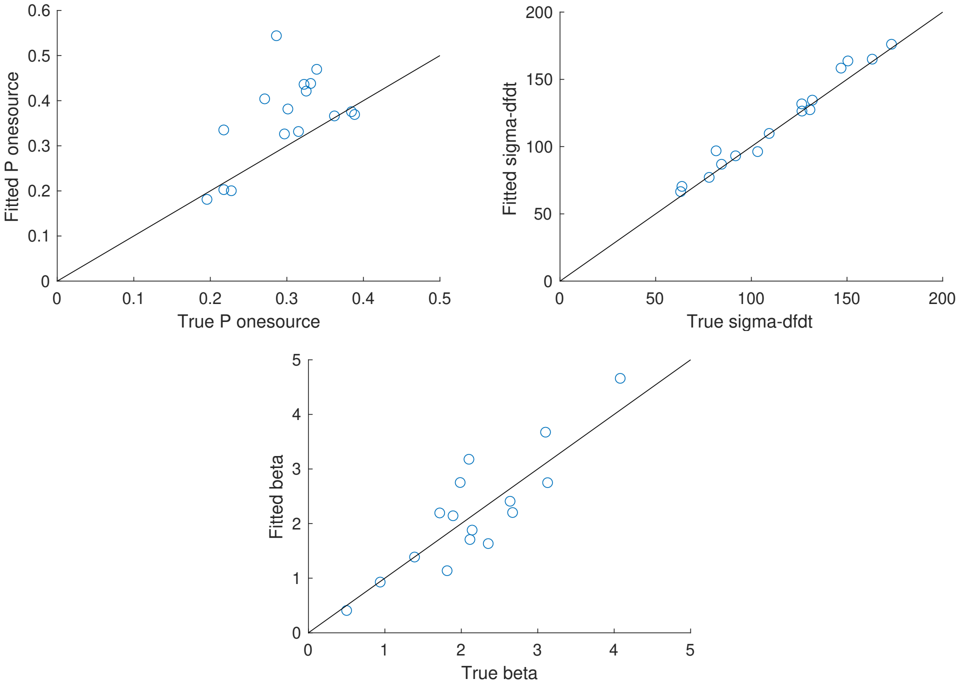


Figure 3: Parameter recovery for Experiment 1. Left) True value for $P(11)$ on horizontal axis, estimated value on the vertical axis. Middle) True value for $\sigma$ on horizontal axis, estimated value on the vertical axis. Right) True value for $\beta$ on horizontal axis, estimated value on the vertical axis.

#### Parameter recovery for Experiment 2 fitting

Methods of parameter recovery were similar to those for Exp 1. Figure 4 (Left) shows that for the $\sigma$ variable the original parameter is quite well recovered. For the $\beta$

parameter in Figure 4 (Right) the recovery is not quite as good (points are further from diagonal) although it is not too surprising given the natural variability of the data when only doing 18 binary choices per condition.

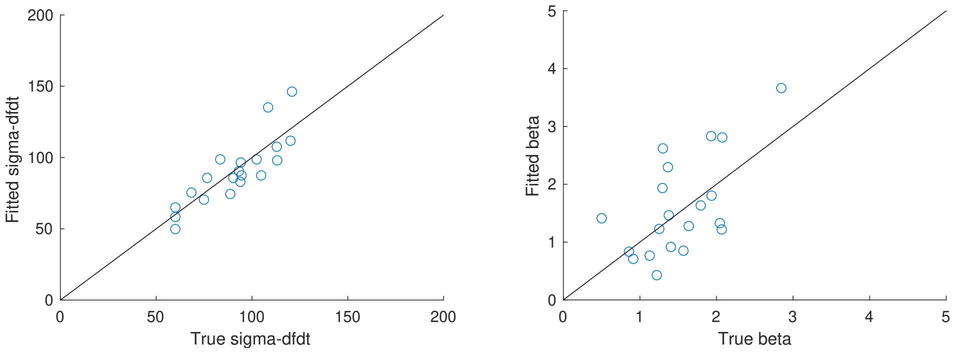


Figure 4: Parameter recovery for Experiment 2. Left) True value for sigma on horizontal axis, estimated value on the vertical axis. Right) True value for beta on horizontal axis, estimated value on the vertical axis.

#### Supporting information - Model recovery

Any attempt at creating a new model faces issues of not just parameter identification and recovery, but also model recovery. A model that (e.g. through error) always assigned a ELBO value of 0 (e.g.) may seem like it is able to explain a lot of data, but is obviously broken. However, this may only be noticable when testing it on data sets where it is known that it should perform poorly (as data were actually generated through a different process).

To ensure that we do not have any issues with the CRP model claiming good model fits on incompatible data, we performed a model recovery exercise. We repeated the methods above (see Parameter recovery above), but for each of the models CRP, alternative A, alternative B, and alternative C. For each of the four models 45 data sets were created (45 x 4 models = 180 datasets). Having generated the simulated data we then fitted each of the four models to each of the data sets (4 x 180 model fits). This allowed us to confirm whether the inferred model, after model fitting and comparison, is indeed the generating 'True' model. Figure 5 shows that while the correct model is generally inferred, occasionally data generated from model CRP (and similarly from model alt B) can be erroneously inferred to be from model alt A. In practice, this means that our model results in the main paper may actually be slightly under estimating the strength of the CRP model.

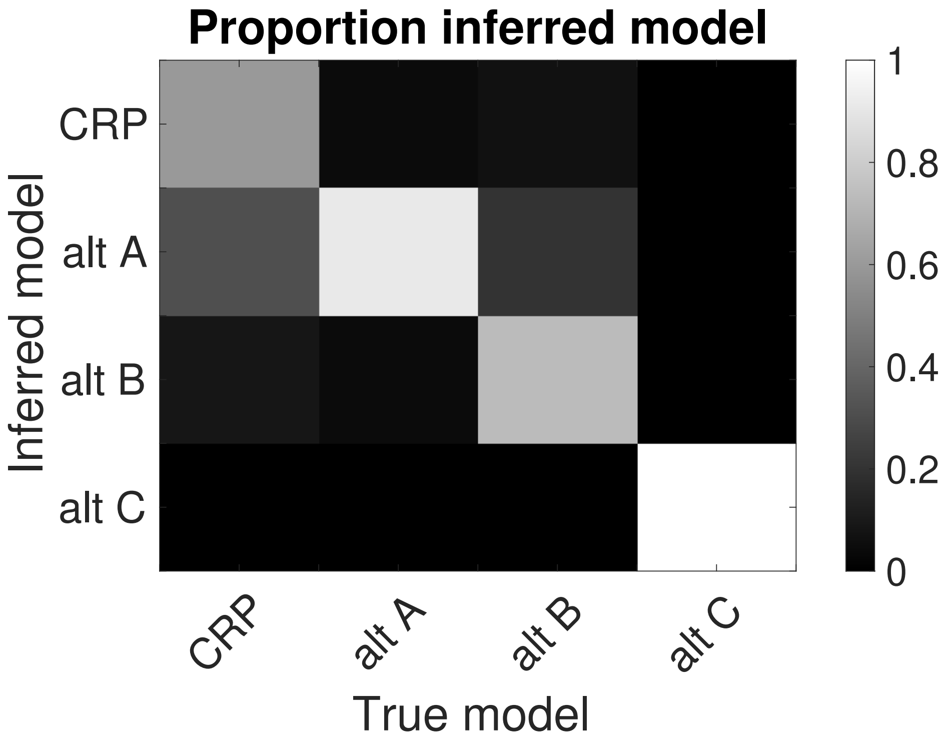


Figure 5: Confusion matrix for model recovery. True generating model on the horizontal axis, while the inferred model after model fitting and comparison is on the vertical axis. In the case of a perfect model recovery the proportion inferred would be zero away from the diagonal.
